# Learning the local landscape of protein structures with convolutional neural networks

**DOI:** 10.1101/2021.08.19.456994

**Authors:** Anastasiya V. Kulikova, Daniel J. Diaz, James M. Loy, Andrew D. Ellington, Claus O. Wilke

## Abstract

The fundamental problem of protein biochemistry is to predict protein structure from amino acid sequence. The inverse problem, predicting either entire sequences or individual mutations that are consistent with a given protein structure, has received much less attention even though it has important applications in both protein engineering and evolutionary biology. Here, we ask whether 3D convolutional neural networks (3D CNNs) can learn the local fitness landscape of protein structure to reliably predict either the wild-type amino acid or the consensus in a multiple sequence alignment from the local structural context surrounding a site of interest. We find that the network can predict wild type with good accuracy, and that network confidence is a reliable measure of whether a given prediction is likely going to be correct or not. Predictions of consensus are less accurate, and are primarily driven by whether or not the consensus matches the wild type. Our work suggests that high-confidence mis-predictions of the wild type may identify sites that are primed for mutation and likely targets for protein engineering.

## 1 Introduction

Proteins are dynamic complex macromolecules that exist as an ensemble of conformational substates (CS) on a fitness landscape [9]. The fitness landscape describes the potential energy of the protein as a function of conformational coordinates [9] and is used to study evolution, to identify proteins with new and useful properties, or to quantify mutability [12]. A protein crystal structure, on the other hand, is a static representation of a protein and is a single point within the energy landscape. This single point is usually a local minimum on the energy landscape, i.e., a highly stable CS that predominates the population and enables crystallization. While a protein crystal structure does not capture the functional and stochastic fluctuations required to model the local energy landscape, sampling thousands of diverse protein crystal structures provides a more global perspective of local energy minima dispersed across the complex energy landscape of all proteins.

The modeling of the protein energy landscape has made significant progress over the last 30 years [12]. Most recently, the application of deep learning to the protein folding problem has shown tremendous success. In particular, Deepmind demonstrated in CASP14 that their deep learning model, alphafold, is able to learn how to generate highly accurate protein structures solely from a protein’s primary sequence and multiple sequence alignment, essentially properly placing a protein sequence in the correct energetic minima [19]. While such great advancements are being made in protein folding, the application of deep learning to the converse problem is lacking: understanding how the structure constrains the amino acids that are allowed at a given site. Prior work using conventional modeling has shown that this is a challenging problem in general [7], and even just predicting whether a site is conserved or variable over evolutionary time is not trivial [23,14,22,16]. Furthermore, the ability to predict which amino acids are “allowed” at a site would limit the sampling of deleterious mutations, accelerating targeted mutagenesis and protein engineering efforts.

Here, we investigate to what extent a 3D self-supervised convolutional neural network (3D CNN) model trained on predicting a masked residue from its local chemical environment–its microenvironment–can predict wild type residues [36,33] and residues in evolutionarily diverged homologs. We show that the CNN model primarily predicts the wild type residue, rather than the residues found in diverged, homologous sequences. Furthermore, we correlate the predicted probability distribution for each residue within the dataset with its observed natural variation in a multiple sequence alignment to examine how much of the natural variation is captured by the 3D CNN model. We assess accuracy as a function of CNN confidence and explore the distribution of amino acids predicted at high confidence. We find that CNN confidence is a good measure of prediction accuracy and that hydrophobic residues are more likely to be predicted with high confidence. Finally, we investigate the impact of microenvironment volume on both the accuracy of predictions and their correlation with natural variation by training several 3D CNN models with different input volumes. Our results demonstrate that the first contact shell plays a crucial role in predicting masked residues from their structural context. Our work may have applications to protein engineering, where sites at which the CNN confidently mis-predicts the resident amino acid may be primed for gain-of-function and hence targets for mutagenesis.

## 2 Methods

### 2.1 Network Architecture

We constructed all convolutional neural network models using tensorflow (v2.4.0) [1]. The network architecture was adopted from the literature [36,33] and consisted of a total of nine layers divided into two blocks: 1) feature extraction and 2) classification. The feature extraction block consisted of six layers: two pairs of 3D convolutional layers followed by a dimension reduction max pooling layer after each pair. The first pair of convolutional layers used filters of size 3 × 3 × 3 and the second pair had filters of size 2 × 2 × 2. Additionally, the Rectified Linearity Unit function (Relu) was applied to the output of each of the four convolutional layers. The final feature maps generated by the feature extraction block had dimensions of size 400 × 3 × 3 × 3. These feature maps were flattened into a 1D vector of size 10, 800 × 1 before being passed to the classification block. The classification block consisted of three fully connected dense layers given dropout rates of 0.5, 0.2, and 0, respectively. Similar to the feature extraction block, the output of the first two dense layers was transformed by the Relu function. To obtain a vector of 20 probability scores representing the network prediction for each of the amino acids, we applied a softmax activation function to the output of the third dense layer. The full list of parameters for each layer in the CNN is provided in Table S1 in Online Resource 1.

### 2.2 Data Generation and Training

To compile the training data, we started with a set of protein crystal structures utilized in previous work [33]. This set provided us with 19,427 distinct protein data bank (PDB) identifiers corresponding to structures with at least a 2.5 Å resolution. Next, we filtered down our dataset by using a 50% sequence similarity threshold at the protein chain level and removing structures where we could not add hydrogen atoms or partial charges in an automated fashion with PDB2PQR (v3.1.0) [5]. Finally, we removed any protein chains that had more than 50% sequence similarity to any structure in the PSICOV dataset [18]. The PSICOV dataset contains 150 extensively studied protein structures and we used it here as a hold-out test dataset to evaluate the CNN models (see below). The final dataset used for training the CNN models consisted of 16,569 chains.

We randomly sampled residues from the protein dataset to create a dataset of microenvironments that reflected the natural abundance of each amino acid. For each protein chain, we sampled at most 100 residues and at most 50% of its residues, whichever number was smaller. We stored the microenvironment dataset as a set of meta-data, recording the specific residues being sampled by their PDB ID, chain ID, residue sequence number, and the wild-type amino acid label. The final microenvironment dataset consisted of 1,455,978 microenvironments. A 90:10 split was utilized for training and validation, respectively. The same training and validation splits were used for all CNN models.

All training instances and labels were generated just-in-time from their microenvironment meta-data, in a batch-wise fashion. With this meta-data, we generate a voxelated representation (4D tensor) of the microenvironment centered at the *α*-carbon of the specified residue, and oriented with respect to the backbone such that the side chain is oriented along the +*z* axis. The voxelated representation had a 1 Å resolution and consisted of 3D space (*x, y, z*) plus 7 auxiliary channels. The auxiliary channels encoded information about the nature of the atom present in a voxel (C, H, O, N, S) as well as partial charge and solvent accessible surface area.

We used PDB2PQR (v3.1.0) [5] to add hydrogen atoms to the protein structure and to calculate the partial charges of each atom in the protein using the PARSE forcefield [34]. We used FreeSASA (v2.0.3) [24] to obtain the solvent accessible surface area of each atom in the protein structure. The spatial dimensions chosen (12 Å, 20 Å, 30 Å, 40 Å) dictated which atoms from the protein structure were included in the voxelated representation of the microenvironments. All atoms of the centered residue were excluded from the voxelated representation.

The CNN models were trained on Radeon Instinct MI-50 accelerator GPUs. All models were trained identically and with the same microenvironment dataset so that model predictions would reflect the impact of modifying the volume of the microenvironment. For each model, we used a batch size of 200 microenvironments and trained for 5 epochs. The loss was calculated with stochastic gradient descent with momentum (0.75) and an adaptive learning rate. The learning rate was initialized at 0.05 and was reduced by half if the validation accuracy did not increase by at least 0.1% every 2,000 batches.

### 2.3 Generating Predictions and Assessing Accuracy

To verify the network performance on an independent dataset, we used the PSICOV dataset [18] as our final test dataset. Each of the PSICOV protein structure PDB files came with multiple sequence alignments (MSAs) ranging from 10 – 60,000 sequences per protein. The PSICOV dataset contained 150 structures, but we discarded 20 of them because either we were unable to add hydrogen atoms or calculate partial charges with PDB2PQR, or the MSAs did not properly align with the sequence in the protein structure. All PSICOV protein structure PDB files and multiple sequence alignments were downloaded from the following archive at Zenodo: https://dx.doi.org/10.5281/zenodo.2552779.

For each of the remaining 130 protein structures, we made boxes around each residue. The size of each box corresponded to the box size with which the network was trained. These boxes were then used as input for the respective trained network, and the network output was a vector of 20 probabilities corresponding to the 20 amino acids for every position in the protein. After generating predictions for each position, the amino acid with the highest probability was identified as the predicted amino acid. These predicted amino acids were compared to the residues from the PSICOV protein structures, referred to as the wild type residues.

To assess the ability of the network to predict the wild type amino acid, we calculated prediction accuracy separately for each protein as the percentage of wild type predictions (i.e., sites where the wild type amino acid is correctly predicted) among the total number of predictions for the protein. To calculate the network accuracy for predicting the amino acid class, individual amino acids were divided into six classes (Table 1) and accuracy calculations previously performed at the level of individual amino acids were repeated at the level of amino acid classes. Finally, to determine whether CNN confidence is a reliable measure of prediction accuracy, we calculated prediction accuracy within probability bins spanning the values of (0–0.2], (0.2–0.4], (0.4–0.6], (0.6–0.8], and (0.8–1.0]. Each bin contained only the positions that were predicted within the specified probability range, and accuracy was again calculated as the percentage of wild type predictions among those positions.

**Table 1.** Amino acids grouped by class.

| class | included amino acids |
| --- | --- |
| aliphatic | Methionine (M), Leucine (L), Inosine (I), Valine (V), Alanine (A) |
| small polar | Cysteine (C), Serine (S), Threonine (T), Asparagine (N), Glutamine (Q) |
| negative | Aspartic Acid (D), Glutamic acid (E) |
| positive | Arginine (R), Lysine (K) |
| aromatic | Histidine (H), Tyrosine (Y), Phenylalanine (F), Tryptophan (W) |
| unique | Proline (P), Glycine (G) |

### 2.4 Comparing Predictions to Sequence Alignments

To compare network predictions to natural sequence alignments, we first found the consensus amino acid for each position in the alignments. We defined the consensus as the amino acid with the highest occurrence at each position in a multiple sequence alignment. In case of ties, we arbitrarily chose one of the tied amino acids as the consensus amino acid at that position. There were only a total 43 ties among ~ 21, 000 positions analyzed.

To determine average sequence divergence of a multiple sequence alignment (MSA) from its reference wild-type sequence, we first calculated the percent similarity of each sequence in the MSA to the wild-type sequence and then averaged over all sequences in the alignment. The percent similarity was calculated by counting the total number of amino acids matching the wild type sequence and dividing by the length of the wild type sequence. In this calculation, gaps in the alignment were treated as mismatches.

To control for sequence divergence, the original PSICOV alignments were divided into five sub-alignment groups based on percent similarity to the wild type sequence. The groups were evenly spaced from 0% similarity to 100% similarity in steps of 20 percentage points. Proteins that did not have 10 or more sequences in their alignments were removed from a group unless they were part of the lowest or highest similarity group (i.e., (0–20%] or (80–100%]).

### 2.5 Site-Specific Variability

For each position in the multiple sequence alignments, we calculated the effective number of amino acids (*n*_eff_) as a measure of site-specific variability. The *n*_eff_ at site *i* is defined as

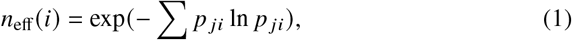

where *p_j i_* is the relative frequency of amino acid *j* at site *i* in the alignment. *n*_eff_ is a number between one and 20, where one indicates that exactly one amino acid is seen at this site and 20 indicates that all twenty amino acids are seen at equal frequencies. Similarly, we calculated an *n*_eff_ corresponding to the neural network predictions, by using the network’s predicted probabilities as the values for *p_j i_* in the above equation.

To compare the variation in the predicted amino acid distributions to the sitespecific variation in alignments, we correlated the vector of *n*_eff_ values obtained from the CNN predictions with the vector of *n*_eff_ values calculated from the MSAs separately for each protein. Similarly, when we binned sequences into five sub-alignments based on percent similarity to the wild type sequence, the *n*_eff_ was calculated for each position in the sub-alignments. For analysis of sub-alignments, we removed all protein structures that did not appear across all 5 similarity bins.

For all correlation coefficients, we assessed statistical significance by calculating *p* values, which we corrected for multiple testing by applying the false-discovery-rate correction [3].

### 2.6 Comparing Amino Acid Distributions

To identify which amino acid types the network is most likely to predict with high confidence, we calculated the frequency with which each amino acid was predicted at a confidence between 80–100%. As a baseline reference, we also calculated the amino acid frequencies in the training data. Finally, to compare the distribution of amino acids that are predicted with high confidence to amino acids that tend to dominate at positions across homologs, we performed similar calculations using the MSAs by calculating the amino acid frequencies at positions where a single residue appears in 80–100% of the sequences in an MSA.

### 2.7 Data availability

Final data analysis and figure production was performed in R [28], making extensive use of the tidyverse family of packages [38]. Analysis scripts and processed data are available on GitHub: https://github.com/akulikova64/CNN_protein_landscape. Trained neural networks and the training set protein chains and microenvironments have been deposited at the Texas Data Repository and are available at: https://doi.org/10.18738/T8/8HJEF9.

## 3 Results

### 3.1 A convolutional neural network predicts wild-type and consensus amino acids with good accuracy

We trained a self-supervised 3D convolutional neural network (3D CNN) to predict the masked amino acid at the center of a chemical environment (microenvironment) extracted from a protein structure. Specifically, the input data to the network consists of a cube with a 1 Å resolution representing all amino-acid atoms surrounding a specific residue (wild-type amino acid), where the specific residue itself has been deleted. Based on this input, the model outputs a discrete probability distribution describing the likelihood of each amino acid being the wild-type amino acid for the given microenvironment. The amino acid with the highest probability is taken as the network’s prediction for the wild-type amino acid at the site. We evaluated the performance of the trained network on an independent dataset of 130 structures (the PSICOV dataset, [18]). All sequences in the training dataset differed by at least 50% from each other and from all sequences in the PSICOV dataset.

We first assessed how well the network could predict a resident amino acid in a protein structure. We refer to the sequence of the PDB structure as the wild type sequence. We found that prediction accuracy for the wild type sequence was generally high, around 60% on average (Figure 1a). In other words, for approximately 60% of all sites in the PSICOV dataset, we could predict the wild-type amino acid from its local chemical environment. We also assessed prediction accuracy at the level of amino-acid classes, where we grouped biochemically similar amino acids into groups (Table 1). We found that our ability to predict amino acid classes was slightly higher than our ability to predict specific amino acids, averaging at approximately 71% (Figure 1a).

**Fig. 1.**
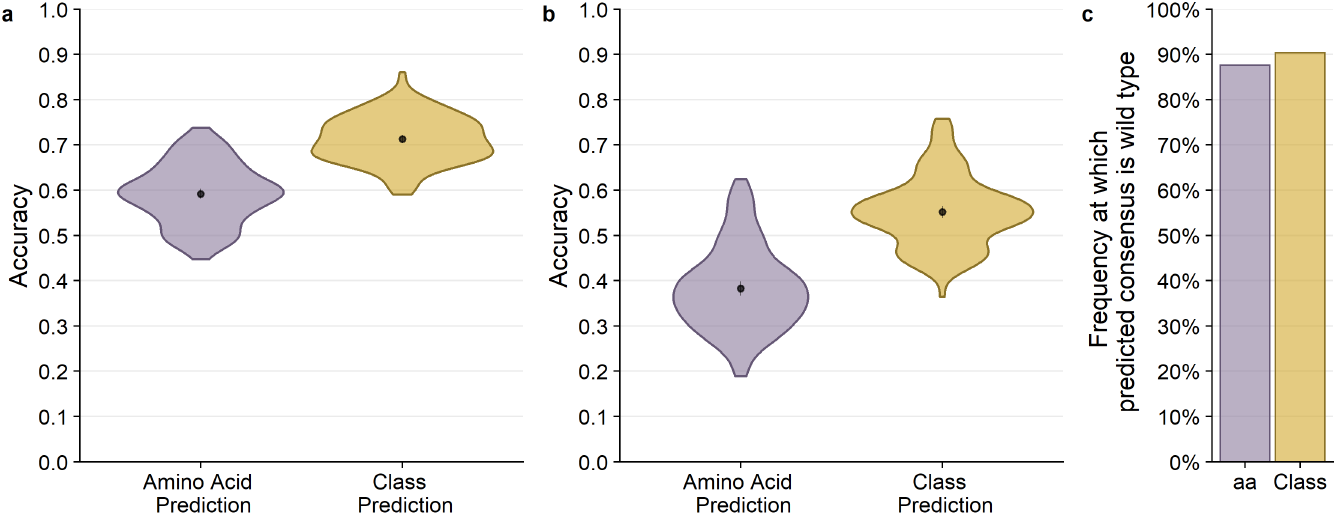
Overall performance of CNN predictions for 20 Å box. The black points and bars represent the means and 95% confidence intervals, respectively. If no bars are visible, the 95% confidence intervals are smaller than the points indicating the location of the means. (a) Prediction accuracy compared to the wild type sequence. Individual amino acids were predicted with a 59.2% mean accuracy and amino acid classes were predicted with 71.3% mean accuracy. (b) Prediction accuracy compared to the alignment consensus. Individual amino acids were predicted with 38.3% mean accuracy and amino acid classes were predicted with 55.2% mean accuracy. (c) The frequency at which the predicted consensus is the wild type. Bars represent the proportion of amino acids (purple) and amino acid classes (yellow) out of all successful consensus predictions that match with the wild type. 86.7% of predicted residues that match the consensus are identical to the wild type and 90.4% of successful consensus amino acid class predictions also match the wild type.

We next asked how well the network could predict the consensus amino acid at a site in a multiple-sequence alignment (MSA). To what extent such a prediction is possible depends on how conserved the microenvironment around a given site is in homologous structures. If biochemical constraints are mostly conserved over evolutionary time, then the network should reliably predict consensus amino acids. By contrast, if the constraints change rapidly as proteins diverge, then the model will poorly predict the consensus amino acid. We found that we could predict the consensus amino acid with approximately 40% accuracy and the consensus class with approximately 55% accuracy (Figure 1b). Notably, nearly all (87%) of the sites where we correctly predicted the consensus amino acid corresponded to sites where the consensus amino acid is identical to the wild-type amino acid (Figure 1c). In other words, the network performs well at predicting the wild-type amino acid, and that extends to the consensus amino acid when it is the same as the wild-type amino acid.

To further elaborate how prediction of the consensus amino acid depends on sequence divergence among homologs, we subdivided alignments into groups with similar sequence similarity to the wild type, calculated the consensus for these similarity groups, and then assessed prediction accuracy for the consensus (Figure 2). As expected, we observed a systematic, linear decline in prediction accuracy with decreasing sequence similarity. For the most diverged sequences (0–20% similarity to wild type), prediction accuracy was below 30%, whereas for the least diverged sequences (80–100% similarity to wild type), accuracy was virtually the same as for predicting wild type (Figure 2a). Results were similar for amino acid classes (Figure 2b). One caveat to this analysis is that homologs were not evenly distributed across the different groups; on average, for each protein, the 80–100% sequence similarity group contained only ~150 sequences whereas the 0–20% similarity group contained ~60,000 sequences (Figure S1 in Online Resource 1). Across all groups, the average number of sequences per protein was ~ 1,000.

**Fig. 2.**
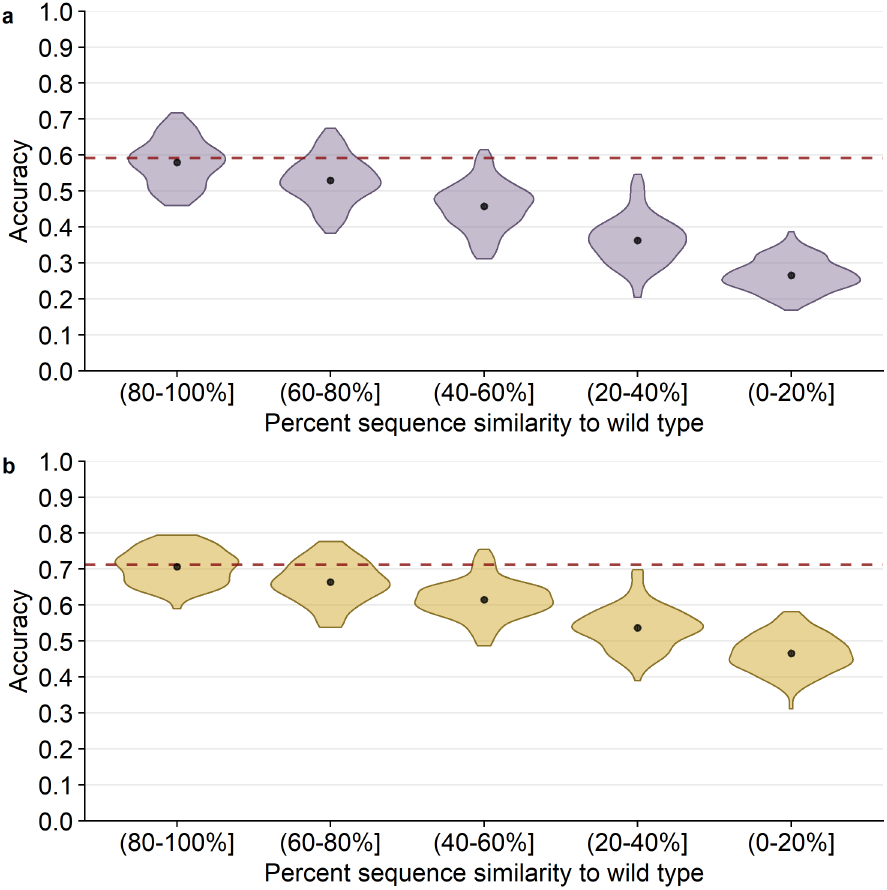
20 Å box CNN predictions of alignment consensus by alignment similarity. The black points and bars represent the means and 95% confidence intervals, respectively. If no bars are visible, the 95% confidence intervals are smaller than the points indicating the location of the means. The percent similarity is the similarity of each alignment sequence to the wild type sequence, the structure of which is used to generate predictions. (a) Percent accuracy of predicting the alignment consensus amino acid. The red dashed line shows the mean accuracy of predicting the wild type residue for comparison (59.2%). (b) Percent accuracy of predicting the alignment consensus class. The red dashed line shows the mean accuracy of predicting the wild type class for comparison (71%).

### 3.2 Network confidence reflects prediction accuracy

Next, we tested whether prediction accuracy was related to CNN confidence. Confidence is defined as the probability with which an amino acid is predicted by the model (i.e., the highest probability in the predicted distribution of all amino acids). We first binned the predicted probabilities into five confidence bins, spanning a range of 0.2 units each, and we then calculated the accuracy of predicting either the wild-type or the consensus amino acid within each confidence bin. We found that the network confidence was an excellent measure of prediction accuracy for wild-type residues (Figure 3a). Mean accuracy almost perfectly matched the CNN confidence for each confidence bin. Similarly, mean accuracy of predicting the consensus residue was consistently at a frequency of 0.1–0.2 below the CNN confidence (Figure 3b). Overall, CNN confidence was a very good measure of prediction accuracy.

**Fig. 3.**
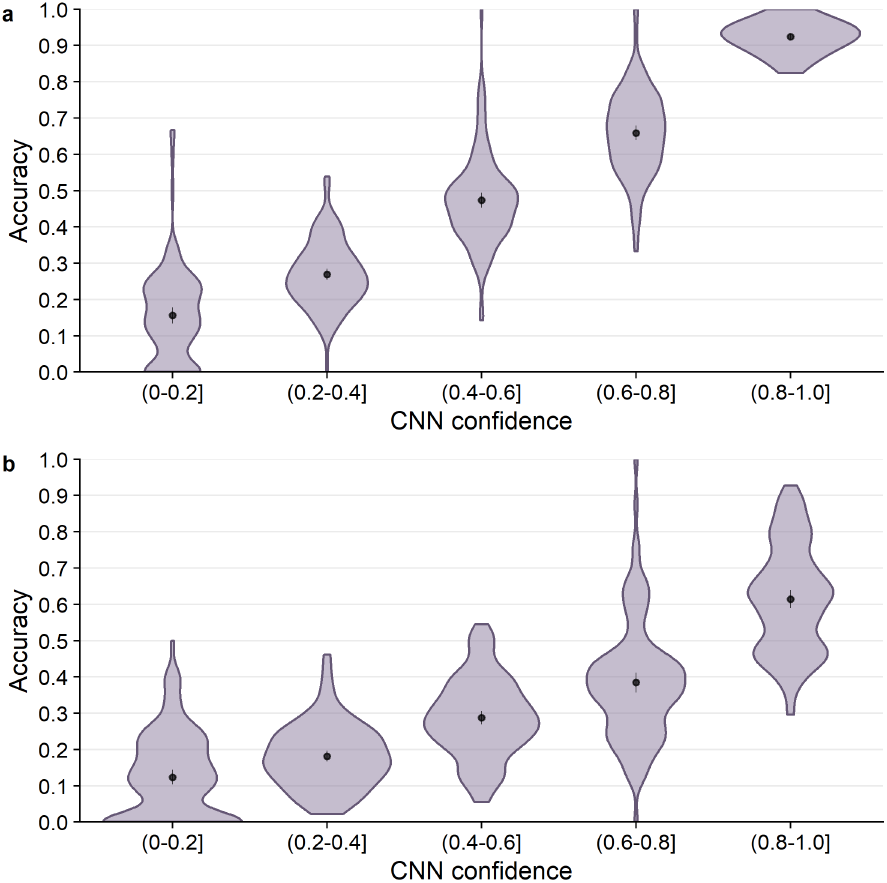
Prediction accuracy as a function of CNN confidence. The black points and bars represent the means and 95% confidence intervals, respectively. If no bars are visible, the 95% confidence intervals are smaller than the points indicating the location of the means. (a) Accuracy of predicting the wild type residue. (b) Accuracy of predicting the consensus residue.

We then wanted to see if high network confidence also implied that all 20 amino acids were predicted with equal likelihood. We selected the positions in the highest predicted probability bin (0.8–1.0] and calculated the frequency with which each amino acid was predicted within this bin. We found that the network predicted the unique and aliphatic (hydrophobic) amino acids with the highest frequencies (Figure 4a). This pattern could be partially explained by the amino acid composition in the training dataset (Figure 4b): The two sets of frequencies were correlated (*r* = 0.658, *p* = 0.002), so approximately 43% of the variation in which amino acids were predicted with high confidence could be explained by the composition of the training dataset. However, by calculating the ratios between these two sets of frequencies, we could confirm that the network trains best on hydrophobic residues (Figure S2 in Online Resource 1). We can speculate that this is because hydrophobic amino acids tend to be found in the core of the protein, where the CNN has more chemical context for generating predictions. By contrast, positive (charged) amino acids were least likely to be predicted with high confidence; these amino acids are more commonly found on the periphery of the protein structure, where the partially empty microenvironment box provides less context for prediction. This finding is consistent with prior studies where a similar network performed poorly on surface residues [33].

**Fig. 4.**
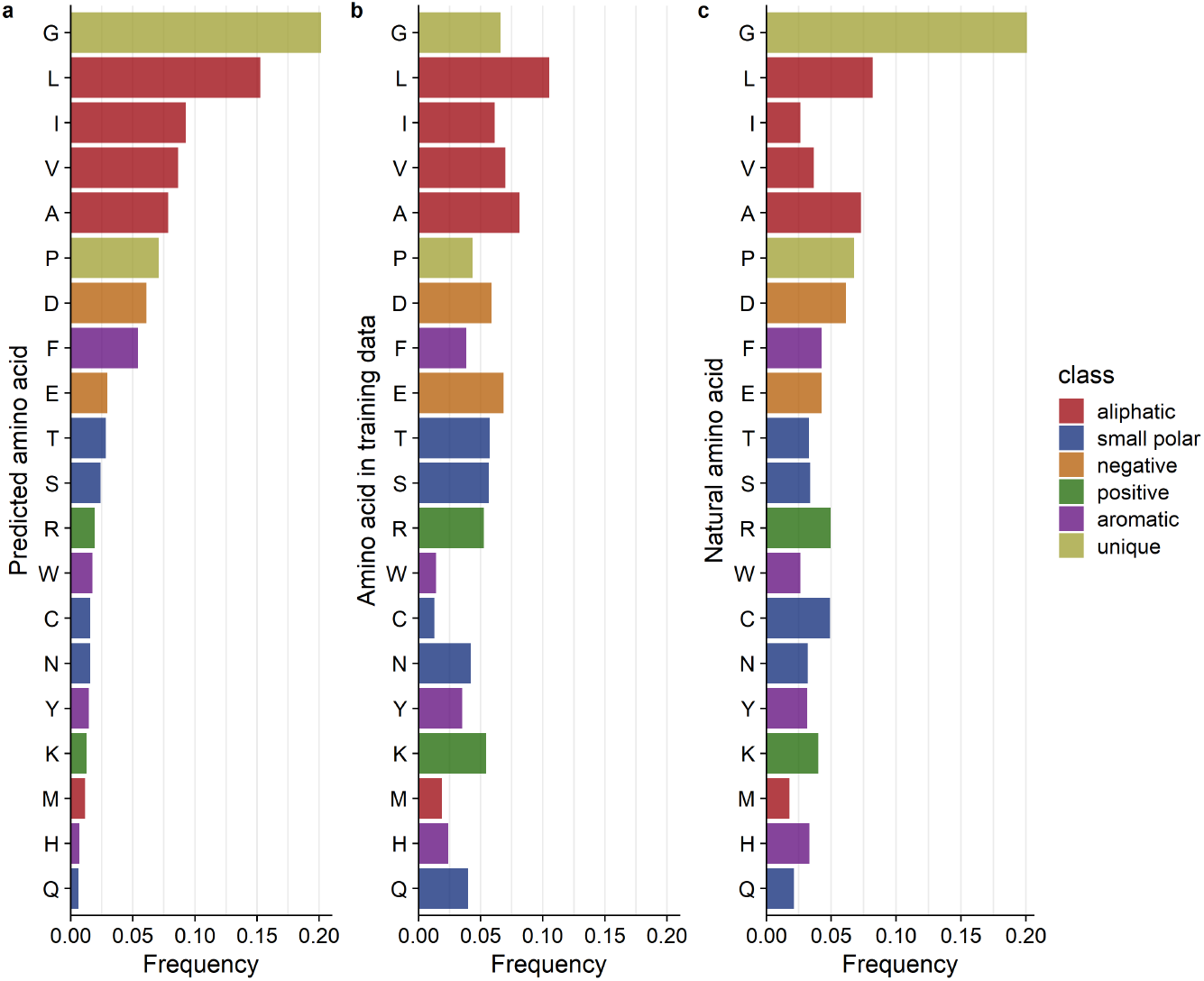
Distributions of amino acid types across predicted, training, and natural datasets. (a) The relative frequency of each amino acid predicted with a confidence of 80–100%. (b) The relative frequency of each amino acid type in the CNN training data. The correlation between predicted and training frequencies was 0.658 (*p* = 0.002) (c) The distribution of highly conserved amino acids. These amino acids are found at frequencies > 80% at individual positions in the MSA. The correlation between predicted and conserved amino acid frequencies was 0.826 (*p* = 0.002).

We also tested whether the same amino acids that tended to receive the highest CNN confidence scores were also the most likely to be conserved in multiple sequence alignments. We extracted positions in the MSAs where the consensus was found in 80– 100% of homologs and calculated the consensus amino acid frequency at these sites (Figure 4c). Interestingly, there was a stronger correlation between the most accurately predicted amino acids and the most conserved ones (*r* = 0.826, *p* = 0.002). The most conserved amino acids tend to be the most easily predicted by the network.

### 3.3 Mis-predictions depend weakly on amino acid biochemistry

We explored the distribution of mis-predictions, predictions that did not match the residue in the wild type structures, by calculating confusion matrices. We calculated confusion matrices at the levels of both individual amino acids and amino acid classes, for both wild-type predictions and consensus predictions (Figure 5). We found some preference of the network to make mis-predictions within the same amino acid class. However, this preference was weak at the level of wild-type predictions (Figure 5a) and only a little more pronounced at the level of consensus predictions (Figure 5c).

**Fig. 5.**
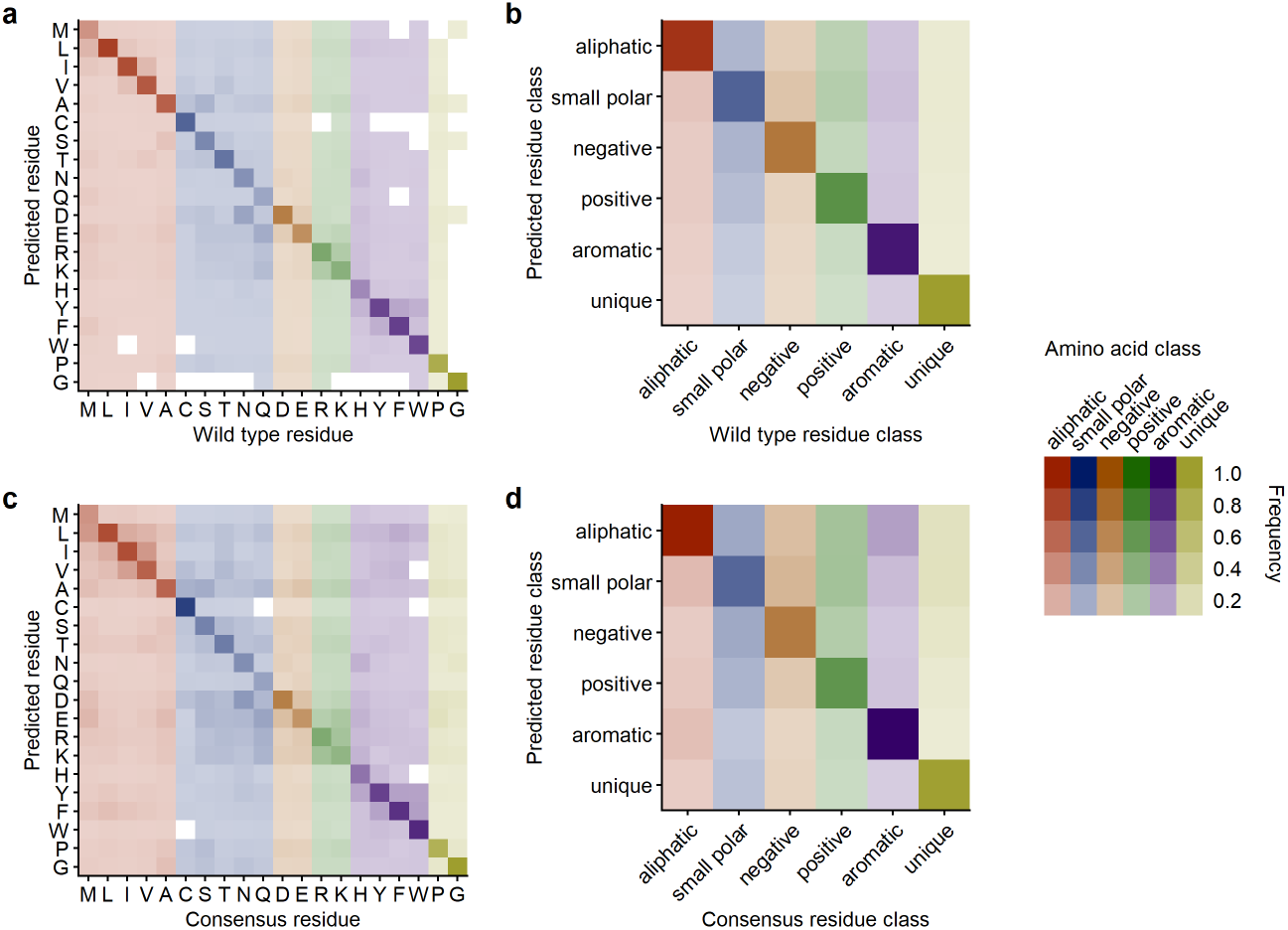
20 Å box CNN prediction frequencies across all amino acids and amino acid classes. (a) Amino acid substitutions relative to wt predictions. (b) Class substitutions relative to wt class predictions. (c) Amino acid substitutions relative to alignment consensus predictions. (d) Class substitutions relative to alignment consensus class predictions.

We then looked only at the network’s mis-predictions and asked if a residue’s predicted probability was related to its abundance at the corresponding position in the MSA. We expected to see that the amino acid substitutions suggested by the CNN model would be found more frequently in nature with increasing CNN confidence. However, we found that the mean natural frequency increased between increasing confidence bins only at a rate of about 2 percentage points. Furthermore, we directly correlated the frequency with which the non-wild-type amino acid is found in homologs with the predicted probability. As expected, the correlation was very weak (*r* = 0.214, *p* < 2 × 10^-16^, Figure S3 in Online Resource 1). We also observed that frequencies vary extensively within each confidence bin and while mis-predicted amino acids sometimes occur at high frequency in natural alignments, in the majority of cases, they occur at near-zero frequency (Figure S3 in Online Resource 1).

### 3.4 Variation in natural alignments correlates weakly with network predictions

As the neural network outputs a probability distribution across all 20 amino acids, we can ask whether the spread in the probability distribution contains any useful information. Specifically, we analyzed whether sites where the network distributes the probability over several amino acids correlate with non-conserved sites in the MSAs and vice-versa. We calculated the correlation of the effective number of amino acids *n*_eff_ at each site for the CNN model predictions and the MSAs, respectively. The effective number is a statistic that ranges from one to 20, where one indicates that only a single amino acid is predicted (CNN) or present (MSA) at the site and 20 indicates that all amino acids are predicted to be equally likely/are present in equal proportions. If the neural network can predict site variability in natural sequence alignments, we would expect the *n*_eff_ calculated from the neural network prediction to correlate with the *n*_eff_ calculated from the MSAs.

We found that correlations varied widely among protein structures but depended only weakly on sequence divergence in the alignment (Figure 6). For some protein structures, we saw significant correlations explaining 10–30% of the variation in the data (correlation coefficients of up to 0.6), while for other protein structures, we saw no significant correlations. Correlations were on average strongest for the three intermediate sequence similarity groups but were not much weaker for other sequence similarity groups.

**Fig. 6.**
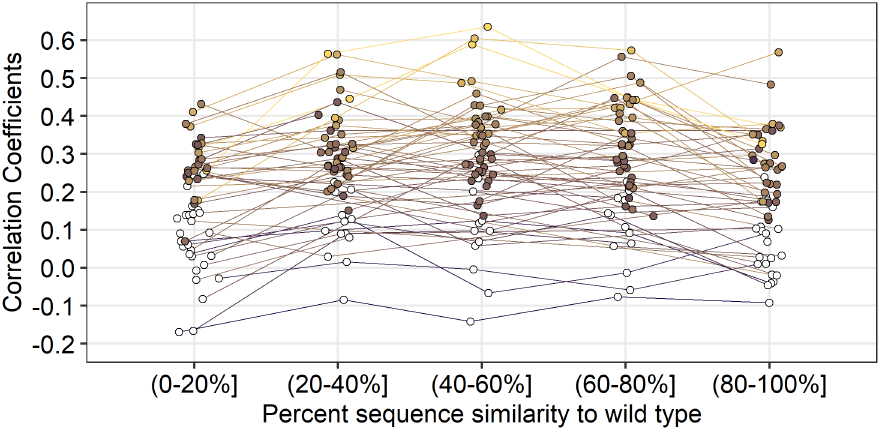
Comparing site-specific variability between CNN predictions and alignments as a function of percent similarity to the wild type. Predictions were generated by the 20 Å box CNN. Variability was calculated as the effective number of amino acids per site (*n*_eff_). Each point represents the correlation coefficient between site-specific predicted variability (*n*_eff_) and alignment variability for a single protein. Colored points represent significant correlations (*p* < 0.05). All *p*-values have been adjusted with the false discovery rate correction. Average significant correlations per similarity group from lowest similarity to highest are 0.270, 0.330, 0.338, 0.335, and 0.286. No significant difference in mean correlation was found between the middle three similarity groups. However, there is a significant increase in mean correlation from the (0–20%] group to the (20–40%] group (*p* = 6 × 10^-07^) and a significant decrease in mean correlation from the (60–80%] group to the (80–100%] group (*p* = 0.0025).

### 3.5 Small microenvironments are sufficient for good network performance

Finally, we assessed whether the volume of the microenvironment had an impact on CNN prediction accuracy. If the CNN model only needs the first contact shell of atoms to accurately generate the probability distribution, then using the smallest possible box that captures the first contact shell should be sufficient. On the other hand, if long-range interactions among amino acids are critical for accurately predicting the probability distribution, then prediction accuracy should increase proportionally with the volume of the microenvironment.

We trained four separate CNN models, using microenvironment volumes of linear size 12 Å, 20 Å, 30 Å, and 40 Å, respectively. Overall, we found that for wild-type amino acid predictions, average accuracy did not change much across box sizes. However, there was a small increase from a mean accuracy of 56.2% to a mean accuracy of 60.6% from the 12 Å box to the 30 Å box (*p* < 1 × 10^-10^)(Figure 7a). The 40 Å box accuracy was lower than the 30 Å box by one percentage point. All differences between consecutive box sizes were between one and three percentage points and statistically significant (*p* < 2 × 10^-16^). Similar patterns were found for the class and consensus predictions (Figure 7a and b).

**Fig. 7.**
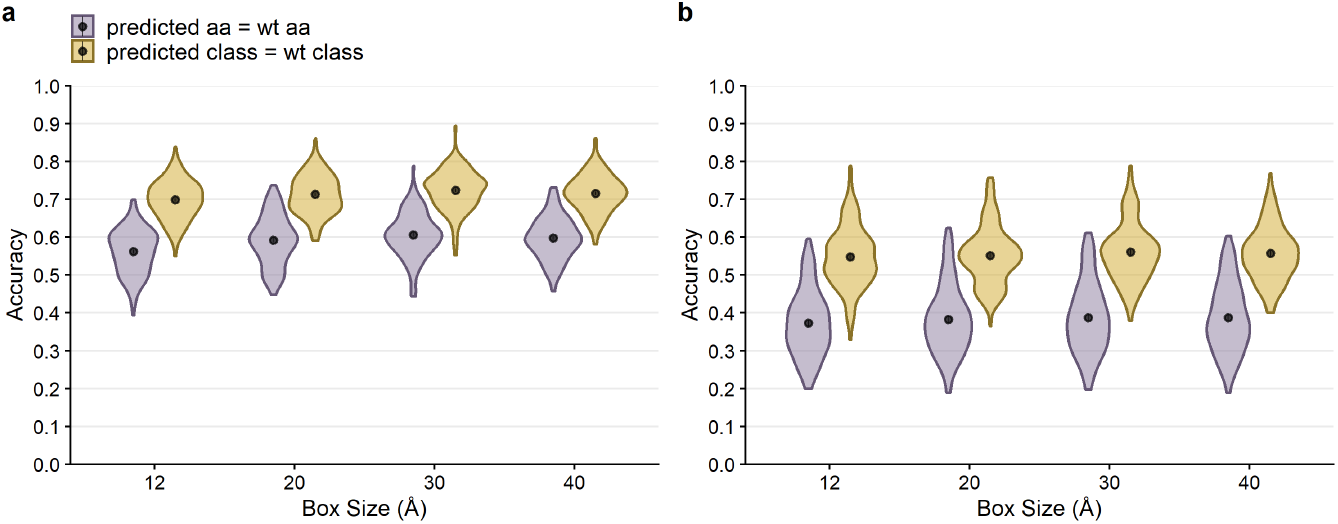
Overall performance of CNN predictions for boxes of size 12 Å, 20 Å, 30 Å and 40 Å. The black points and bars represent the means and 95% confidence intervals, respectively. If no bars are visible, the 95% confidence intervals are smaller than the points indicating the location of the means. All *p*-values have been adjusted with the false discovery rate correction. (a) Prediction accuracy compared to the wild type sequence. The average amino acid accuracies for box sizes of 12, 20, 30, and 40 Å were 0.562, 0.592, 0.606, and 0.598, respectively, and the average class accuracies were 0.698, 0.713, 0.724, and 0.716, respectively. For amino acid predictions (purple), the differences in mean accuracy between all consecutive box sizes were significant (*p* < 9 × 10^-05^). (b) Prediction accuracy compared to the alignment consensus. The average amino acid accuracies for box sizes of 12, 20, 30, and 40 Å were 0.373, 0.383, 0.388, and 0.387, respectively, and the average class accuracies were 0.548, 0.552, 0.561, and 0.557, respectively. For consensus amino acid predictions (purple), only the difference in accuracy between the 12 Å box predictions and the other three box sizes was found to be significant (*p* < 0.003). Accuracy is slightly lower for the 12 Å box than for the other three box sizes.

We also assessed whether microenvironment volume had an effect on the network’s ability to learn natural variation. Because the (40–60%] similarity group had the highest correlation between predicted and natural *n*_eff_ for the 20 Å network, we restricted our analysis to alignments in this similarity group. We repeated the analysis from Figure 6 and found that the correlations between predicted and observed site variability also did not appear to vary much with microenvironment volume (Figure S4 in Online Resource 1). However, the highest correlations were observed for the smallest two microenvironment volumes. In aggregate, these results suggest that the atoms in the first contact shell hold most of the information the CNN model is using to generate the probability distribution, and additional atoms present in larger microenvironment volumes are not contributing much additional information.

## 4 Discussion

We have examined the ability of a Convolutional Neural Network (CNN) to predict resident amino acids in a protein structure from their local microenvironment. The self-supervised learning task and data engineering methodologies were adopted from prior work [36,33]. We have found that the CNN model primarily predicts the wild type residue and that its confidence is a reliable estimate of prediction accuracy. Looking at amino acids predicted with highest confidence, we have found little association with their conservation in homologous sequences. Finally, by training four separate CNNs using different microenvironment volumes as input data, we have learned that the network primarily uses the first contact shell; larger microenvironments do not add much additional information to the prediction task.

Our findings reinforce the notion that amino acids are entrenched in their local biochemical environment [31,11], and that the constraints acting on an amino acid in a protein tend to change over time as the protein evolves. We can predict the resident amino acid at a site from its structural surroundings with approximately 60% accuracy — far better than random chance. Notably, though, 40% of the time we predict the wrong amino acid, and 30% of the time we predict the wrong amino-acid class. There are two reasons why our predictions may be wrong, and we expect that our dataset contains both cases: (i) the neural network is making an incorrect prediction; (ii) the neural network prediction is correct, and the resident amino acid is actually inconsistent with its current chemical surroundings. The second scenario provides an opportunity for protein engineering, as it would point out sites that are primed for mutation. Theoretical arguments and simulations suggest such sites must exist [31, 11], and future work will have to determine whether we can reliably identify such sites and make use of them in protein engineering applications. A small pilot study suggests that this indeed is the case [33].

Predictions at the alignment level (i.e., predicting the consensus amino acid in a multiple sequence alignment) are substantially less accurate — around 40% on average — though still much better than random chance. This finding again reinforces that sites are not independent in protein structures and that the constraints on individual sites slowly change over time [27]. Our analysis of prediction accuracy as a function of sequence similarity provides insight into the extent to which local environments change as mutations accumulate. When going from the highest sequence similarity group (80–100%) to the lowest sequence similarity group (0–20%), prediction accuracy drops approximately by half. Thus, even if we replace nearly every amino acid in a protein, the network prediction accuracy declines only by about 50%. This shows that biochemical environments change only slowly and can be conserved even at extreme levels of sequence divergence. Similar observations have been made previously in the context of protein–protein binding interfaces [20,35]. Additionally, it would be interesting to see if feeding the CNN models molecular dynamic simulations of the microenvironments could enable a better assessment the biochemical environment around an amino acid and potentially improve how well the CNN model predictions recapitulate natural variation.

Our findings also confirm previously leveled criticism [10] against widely used exchangeability matrices such as BLOSUM, WAG, or LG [13,37,21]. These matrices assume that we can meaningfully define exchangeability scores that predict how easily we can replace an amino acid with any other amino acid at any site in a protein, and they predict that virtually any amino acid can be replaced with any other. The problem with this prediction is that while it may be true when averaging over millions of sites and many thousand proteins, exchangeability predictions are virtually useless at any given site of interest. In practice, at any given site of interest, only a small number (~ 5 on average) of amino acids are allowed [10,8,17], and mutations to other amino acids will rarely if ever be observed. Similarly, here we have seen that the local biochemical environment, which is entirely ignored by exchangeability matrices, provides a strong constraint on the resident amino acid, and it predicts the resident amino acid in at least 60% of all cases. We believe that exchangeability models based on local amino acid preferences and/or local biochemical environments are much more useful than global exchangeability models that treat every single site the same.

Looking only at residues predicted with an accuracy of 80% and above, we have not seen equal prediction frequencies across all 20 amino acids. In part, this is driven by natural amino acid abundance. For example, hydrophobic/non-polar amino acids, which include aliphatic and certain aromatic amino acids, comprise a larger portion of the protein, as they are most abundant in the protein core [6]. Consequently, they are also more frequent in our training dataset. However, we have found prediction accuracy of hydrophobic/non-polar amino acids to be higher than expected given their frequencies in the training dataset. One possibility that would explain this finding is that hydrophobic residues, because they tend to be located in the core, are more commonly found in microenvironments that are entirely filled instead of partially empty, providing the CNN more context to make accurate predictions.

We were surprised to find good prediction accuracy with small microenvironment volumes, as long-range interactions in proteins are well documented [15,2,25,32]. This reinforces the notion that long-range interactions act via percolation through the structure. Because the CNN model’s predictions remain localized to a small microenvironment, the model does not generalize it’s predictions to more distant contacts. Consequently, we cannot expect the model to successfully predict a substitution at one site without being required to predict compensatory substitutions at another site [26]. Furthermore, it would be inaccurate to rule out any single network-suggested substitutions if they do not immediately produce a functional protein; additional compensatory substitutions may be required. Unfortunately, the current model can not account for combinations or groups of substitutions.

Our research aims to shed light on the applicability of machine learning for predicting a protein’s fitness landscape. Due to current advancements in next-generation sequencing and the ever growing collection of digital, chemical, and biological data, computational models are becoming increasingly popular for learning the relationship between protein sequence and structure. For example, Hidden Markov Models are being developed for reconstructing protein fitness landscapes using families of homologs [4]. Neural network-based approaches, including Transformers and Generative Adversarial Networks (GANs), have also become increasingly widely used for the prediction and simulation of functional sequence space [39,30,29]. Recently, the deep neural network-based model developed by Deepmind, alphafold, has shown state of the art success in predicting the global protein structure from sequence [19]. Here, we have addressed the converse problem, where Convolutional Neural Networks (CNNs) are trained to learn and predict a protein’s sequence from its local structure [36,33]. Going forward, we expect that the primary application for this converse problem is going to be protein engineering. CNNs such as the one studied here can be used to identify sites that are primed for mutation because the resident amino acid is not consistent with its microenvironment [36,33]. Future work will have to determine whether best results are obtained if the network is trained on wild-type residues, as was done here, or whether training the network on natural variation will yield better candidates for protein engineering.

## Supporting information

Online Resource 1

## Declarations

### Availability of data, materials, and code

Analysis scripts and processed data are available on GitHub: https://github.com/akulikova64/CNN_protein_landscape. Trained neural networks and the training set protein chains and microenvironments have been deposited at the Texas Data Repository and are available at: https://doi.org/10.18738/T8/8HJEF9.

### Authors’ contributions

All authors contributed to the study conception and design. Material preparation, data collection and analysis were performed by A. V. Kulikova, D. J. Diaz, and J. M. Loy. The first draft of the manuscript was written by A. V. Kulikova and all authors commented on previous versions of the manuscript. All authors read and approved the final manuscript.

## References

1. Abadi, M., Agarwal, A., Barham, P., E., B., Chen, Z., Citro, C., Corrado, G.S., Davis, A., Dean, J., Devin, M., Ghemawat, S., Goodfellow, I., Harp, A., Irving, G., Isard, M., Jozefowicz, R., Jia, Y., Kaiser, L., Kudlur, M., Levenberg, J., Mané, D., Schuster, M., Monga, R., Moore, S., Murray, D., Olah, C., Shlens, J., Steiner, B., Sutskever, I., Talwar, K., Tucker, P., Vanhoucke, V., Vasudevan, V., Viégas, F., Vinyals, O., Warden, P., Wattenberg, M., Wicke, M., Yu, Y., Zheng, X.: Tensorflow: Large-scale machine learning on heterogeneous systems (2015). Software available from:https://www.tensorflow.org/

2. Abriata, L.A., Bovigny, C., Dal Peraro, M.: Detection and sequence/structure mapping of biophysical constraints to protein variation in saturated mutational libraries and protein sequence alignments with a dedicated server. BMC Bioinf. 17, 242 (2016)

3. Benjamini, Y., Hochberg, Y.: Controlling the false discovery rate: A practical and powerful approach to multiple testing. J R Stat Soc Series B Stat Methodol J R STAT SOC B 57, 289–300 (1995)

4. Bisardi, M., Rodriguez-Rivas, J., Zamponi, F., Weigt, M.: Modeling sequence-space exploration and emergence of epistatic signals in protein evolution (2021)

5. Dolinsky, T.J., Czodrowski, P., Li, H., Nielsen, J.E., Jensen, J.H., Klebe, G., Baker, N.A.: PDB2PQR: expanding and upgrading automated preparation of biomolecular structures for molecular simulations. Nucleic Acids Research 35, W522–W525 (2007)

6. Dyson, H.J., Wright, P.E., Scheraga, H.A.: The role of hydrophobic interactions in initiation and propagation of protein folding. Proc. Natl. Acad. Sci. U.S.A. 103(35), 13057–13061 (2006)

7. Echave, J., Spielman, S.J., Wilke, C.O.: Causes of evolutionary rate variation among protein sites. Nature Rev. Genet. 17, 109–121 (2016)

8. Echave, J., Wilke, C.O.: Biophysical models of protein evolution: understanding the patterns of evolutionary sequence divergence. Annu. Rev. Biophys. 46, 85–103 (2017)

9. Frauenfelder, H., Sligar, S.G., Wolynes, P.G.: The energy landscapes and motions of proteins. Science 254, 1598–1603 (1991)

10. Goldstein, R.A., Pollock, D.D.: The tangled bank of amino acids. Protein Sci. 25, 1354–1362 (2016)

11. Goldstein, R.A., Pollock, D.D.: Sequence entropy of folding and the absolute rate of amino acid substitutions. Nature Ecol. Evol. 1, 1923–1930 (2017)

12. Hartman, E.C., Tullman-Ercek, D.: Learning from protein fitness landscapes: a review of mutability, epistasis, and evolution. Curr. Opin. Syst. Biol. 14, 25–31 (2019)

13. Henikoff, S., Henikoff, J.G.: Amino acid substitution matrices from protein blocks. Proc. Natl. Acad. Sci. USA 89, 10915–10919 (1992)

14. Huang, T.T., d. V. Marcos, M.L., Hwang, J.K., Echave, J.: A mechanistic stress model of protein evolution accounts for site-specific evolutionary rates and their relationship with packing density and flexibility. BMC Evol. Biol. 14(2014)

15. Jack, B.R., Meyer, A.G., Echave, J., Wilke, C.O.: Functional sites induce long-range evolutionary constraints in enzymes. PLOS Biol. 14, 1–23 (2016)

16. Jiang, Q., A. I. Teufel E. L. Jackson, C.O., Wilke: Beyond thermodynamic constraints: Evolutionary sampling generates realistic protein sequence variation. Genetics 208, 1387–1395 (2018)

17. Johnson, M.M., Wilke, C.O.: Site-specific amino acid distributions follow a universal shape. J. Mol. Evol. 88, 731–741 (2020)

18. Jones, D.T., Buchan, D.W.A., Cozzetto, D., Pontil, M.: PSICOV: precise structural contact prediction using sparse inverse covariance estimation on large multiple sequence alignments. Bioinformatics 28, 184–190 (2011)

19. Jumper, J., Evans, R., Pritzel, A., Green, T., Figurnov, M., Ronneberger, O., Tunyasuvunakool, K., Bates, R., Žídek, A., Potapenko, A., Bridgland, A., Meyer, C., Kohl, S.A.A., Ballard, A.J., Cowie, A., Romera-Paredes, B., Nikolov, S., Jain, R., Adler, J., Back, T., Petersen, S., Reiman, D., Clancy, E., Zielinski, M., Steinegger, M., Pacholska, M., Berghammer, T., Bodenstein, S., Silver, D., Vinyals, O., Senior, A.W., Kavukcuoglu, K., Kohli, P., Hassabis, D.: Highly accurate protein structure prediction with AlphaFold. Nature (2021). DOI 10.1038/s41586-021-03819-2

20. Kachroo, A.H., Laurent, J.M., Yellman, C.M., Meyer, A.G., Wilke, C.O., Marcotte, E.M.: Systematic humanization of yeast genes reveals conserved functions and genetic modularity. Science 348, 921–925 (2015)

21. Le, S.Q., Gascuel, O.: An improved general amino acid replacement matrix. Mol. Biol. Evol. 25, 1307–1320 (2008)

22. Marcos, M.L., Echave, J.: Too packed to change: side-chain packing and site-specific substitution rates in protein evolution. PeerJ 3, e911 (2015)

23. Mirny, L.A., Shakhnovich, E.I.: Universally conserved positions in protein folds: reading evolutionary signals about stability, folding kinetics and function. J. Mol. Biol. 291, 177–196 (1999)

24. Mitternacht, S.: FreeSASA: an open source C library for solvent accessible surface area calculations [version 1; peer review: 2 approved]. F1000Research 5, 189 (2016)

25. Nelson, E.D., Grishin, N.V.: Long-range epistasis mediated by structural change in a model of ligand binding proteins. PLoS ONE 11, e0166739 (2016)

26. Pokusaeva, V.O., Usmanova, D.R., Putintseva, E.V., Espinar, L., Sarkisyan, K.S., Mishin, A.S., Bogatyreva, N.S., Ivankov, D.N., Akopyan, A.V., Avvakumov, S.Y., Povolotskaya, I.S., Filion, G.J., Carey, L.B., Kondrashov, F.A.: An experimental assay of the interactions of amino acids from orthologous sequences shaping a complex fitness landscape. PLOS Genet. 15, 1–30 (2019)

27. Pollock, D.D., Thiltgen, G., Goldstein, R.A.: Amino acid coevolution induces an evolutionary Stokes shift. Proc. Natl. Acad. Sci. USA 109, E1352–E1359 (2012)

28. R Core Team: R: a language and environment for statistical computing. R Foundation for Statistical Computing, Vienna, Austria (2019)

29. Repecka, D., Jauniskis, V., Karpus, L., Rembeza, E., Rokaitis, I., Zrimec, J., Poviloniene, S., Laurynenas, A., Viknander, S., Abuajwa, W., Savolainen, O., Meskys, R., Engqvist, M.K.M., Zelezniak, A.: Expanding functional protein sequence spaces using generative adversarial networks. Nat. Mach. Intell. 3, 324–333 (2021)

30. Rives, A., Meier, J., Sercu, T., Goyal, S., Lin, Z., Liu, J., Guo, D., Ott, M., Zitnick, C.L., Ma, J., Fergus, R.: Biological structure and function emerge from scaling unsupervised learning to 250 million protein sequences. Proc. Natl. Acad. Sci. U.S.A. 118(15) (2021)

31. Shah, P., McCandlish, D.M., Plotkin, J.B.: Contingency and entrenchment in protein evolution under purifying selection. Proc. Natl. Acad. Sci. USA 112, E3226–E3235 (2015)

32. Sharir-Ivry, A., Xia, Y.: Nature of long-range evolutionary constraint in enzymes: insights from comparison to pseudoenzymes with similar structures. Mol. Biol. Evol. 35, 2597–2606 (2018)

33. Shroff, R., Cole, A.W., Diaz, D.J., Morrow, B.R., Donnell, I., Gollihar, J., Ellington, A.D., Thyer, R.: Discovery of novel gain-of-function mutations guided by structure-based deep learning. ACS Synth. Biol. 9, 2927–2935 (2020)

34. Sitkoff, D., Sharp, K.A., Honig, B.: Accurate calculation of hydration free energies using macroscopic solvent models. J. Phys. Chem. 98, 1978–1988 (1994)

35. Teufel, A.I., Johnson, M.M., Laurent, J.M., Kachroo, A.H., Marcotte, E.M., Wilke, C.O.: The many nuanced evolutionary consequences of duplicated genes. Mol. Biol. Evol. 36, 304–314 (2019)

36. Torng, W., Altman, R.B.: 3D deep convolutional neural networks for amino acid environment similarity analysis. BMC Bioinf. 18, 302 (2017)

37. Whelan, S., Goldman, N.: A general empirical model of protein evolution derived from multiple protein families using a maximum-likelihood approach. Mol. Biol. Evol. 18, 691–699 (2001)

38. Wickham, H., Averick, M., Bryan, J., Chang, W., D’Agostino McGowan, L., François, R., Grolemund, G., Hayes, A., Henry, L., Hester, J., Kuhn, M., Lin Pedersen, T., Miller, E., Milton Bache, S., Müller, K., Ooms, J., Robinson, D., Paige Seidel, D., Spinu, V., Takahashi, K., Vaughan, D., Wilke, C., Woo, K., Yutani, H.: Welcome to the tidyverse. J. Open Source Softw. 4, 1686 (2019)

39. Xu, Y., Verma, D., Sheridan, R.P., Liaw, A., Ma, J., Marshall, N.M., McIntosh, J., Sherer, E.C., Svetnik, V., Johnston, J.M.: Deep dive into machine learning models for protein engineering. J. Chem. Inf. Model. 60, 2773–2790 (2020)

