## Supplementary material for "Learning the local landscape of protein structures with convolutional neural networks": Online Resource 1

### Supplementary Materials for Kulikova et al., Learning the local landscape of protein structures with convolutional neural networks

Table S1: Full list of parameters for each layer in the CNN. The feature extraction block consists of layers 1–6, while layers 7–9 are part of the classification block. All convolutional or max-pooling layers are 3-dimensional.

| layer | layer type | parameters |
|---|---|---|
| 1 | convolutional | filters = 100, kernel = $(3 \times 3 \times 3)$, stride = 1, activation = ReLu, padding = False |
| 2 | convolutional | filters = 200, kernel = $(3 \times 3 \times 3)$, stride = 1, activation = ReLu, padding = False |
| 3 | max-pooling | no parameters |
| 4 | convolutional | filters = 200, kernel = $(2 \times 2 \times 2)$, stride = 1, activation = ReLu, padding = False) |
| 5 | convolutional | filters = 400, kernel = $(2 \times 2 \times 2)$, stride = 1, activation = ReLu, padding = False) |
| 6 | max-pooling | no parameters |
| 7 | dense | neurons = 1000, dropout = 0.5, activation = ReLu |
| 8 | dense | neurons = 100, dropout = 0.2, activation = ReLu |
| 9 | dense | neurons = 20, activation = softmax |

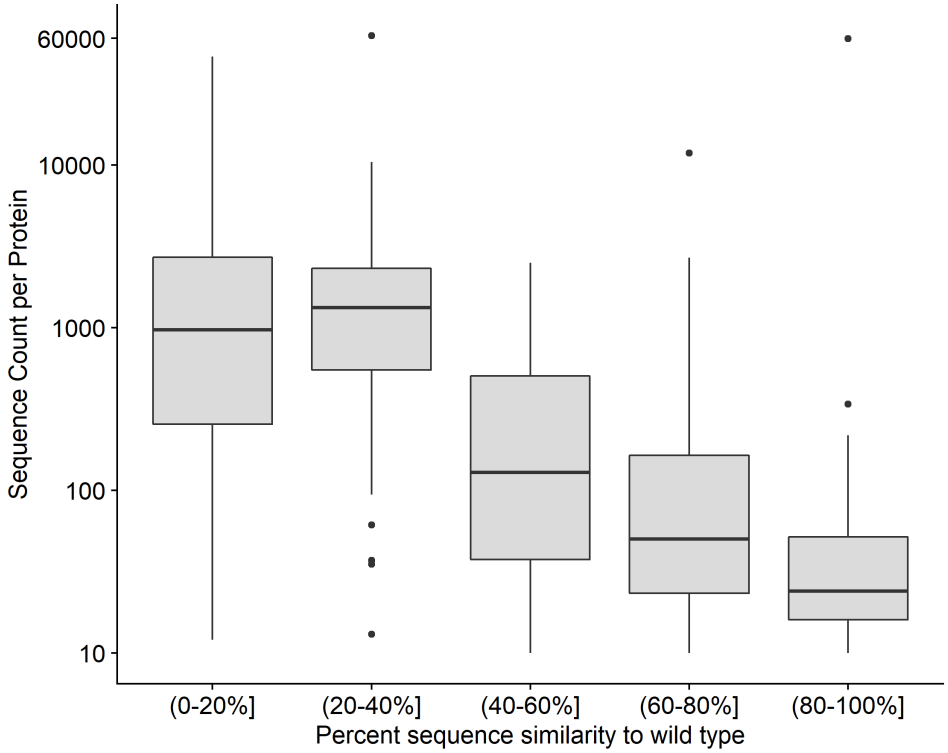


Figure S1: Number of sequences per alignment for each percent similarity group. The lowest number of sequences per alignment is found in the least diverged group. The two most diverged groups range from 10 to 60,000 sequences per alignment.

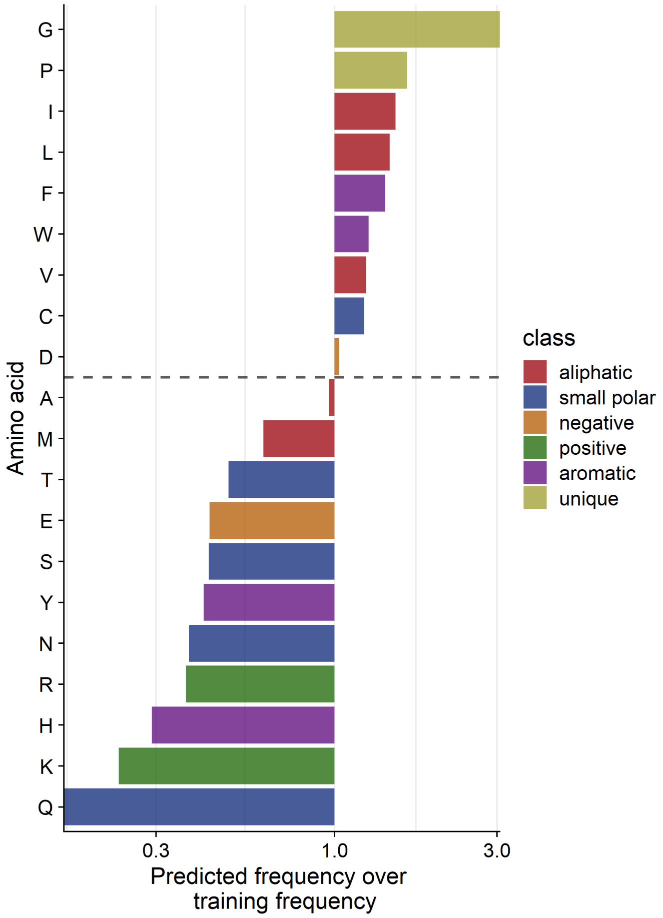


Figure S2: The frequency of the amino acid confidently predicted over the frequency the amino acid found in the training dataset. The grey dashed line divides amino acids with ratios greater or less than 1. Ratios greater than 1 indicate that the neural network confidently predicts those amino acids more often than their frequency in training data. A ratio less than 1 indicates that the amino acid is found in the training data a higher frequency than predicted by the network.

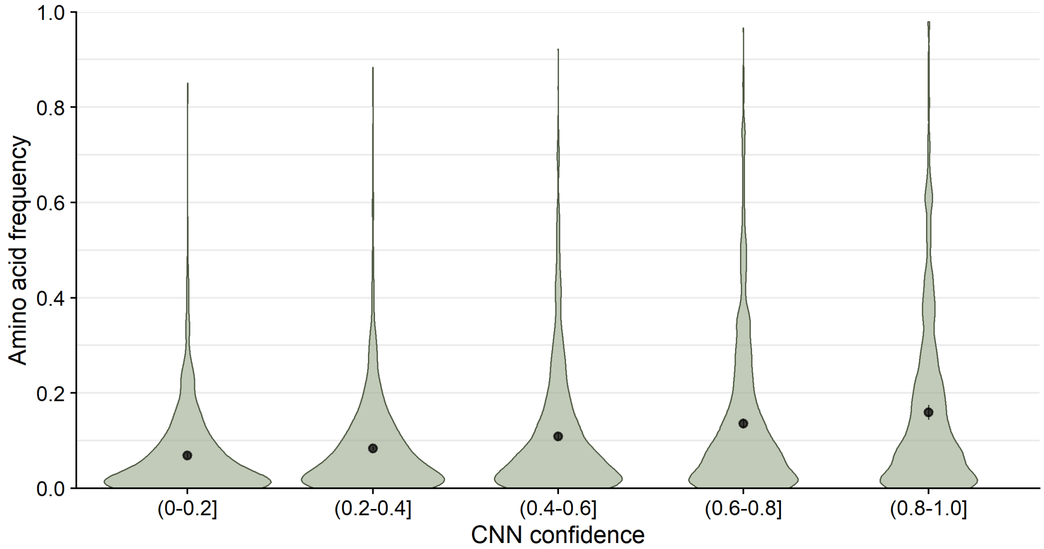


Figure S3: Natural Frequency of the predicted amino acid when it is not the wild type. The black points and bars represent the means and 95% confidence intervals, respectively. If no bars are visible, the 95% confidence intervals are smaller than the points indicating the location of the means. Mean frequencies for each CNN confidence bin were 0.069, 0.084, 0.109, 0.136 and 0.159, in order of increasing confidence. The number of positions per confidence bin were 1070, 3141, 1711, 924, and 510, respectively.

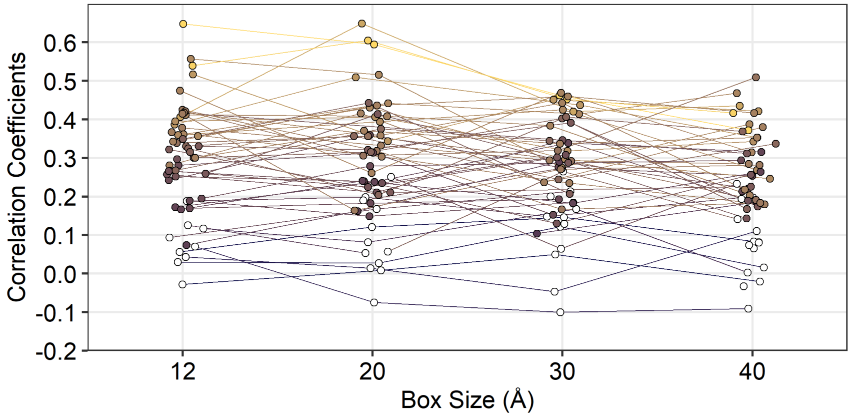


Figure S4: Comparing site-specific variability between boxes of size 12 Å, 20 Å, 30 Å and 40 Å for the alignment with 40–60% similarity. Variability was calculated as the effective number of amino acids per site ($n_{\text{eff}}$). Each point represents the correlation coefficient between site-specific predicted variability ($n_{\text{eff}}$) and alignment variability for a single protein. Colored points represent significant correlations ($p < 0.05$). All $p$-values have been adjusted with the false discovery rate correction. Mean correlations were 0.291, 0.285, 0.266 and 0.235, for the 12, 20, 30 and 40 Å boxes, respectively. A paired t-test showed significant difference for 5 out of 6 pairs of box sizes ($p < 0.03$). Only the correlations between the 12 Å and 20 Å box did not show significant difference ($p = 0.4402$).
